# Epidemiological inference at the threshold of data availability: an influenza A(H1N2)v spillover event in the United Kingdom

**DOI:** 10.1101/2024.03.11.584378

**Authors:** John A. Fozard, Emma C. Thomson, Christopher J. R. Illingworth

## Abstract

Viruses which infect animals regularly spill over into the human population, but individual events may lead to anything from a single case to a novel pandemic. Rapidly gaining an understanding of a spillover event is critical to calibrating a public health response. We here propose a novel method, using likelihood free rejection sampling to evaluate the properties of an outbreak of swine-origin influenza A(H1N2)v in the United Kingdom, detected in November 2023. From the limited data available we generate historical estimates of the probability that the outbreak had died out in the days following the detection of the first case. Our method suggests that the outbreak could have been said to be over with 95% certainty between 19 and 29 days after the first case was detected, depending upon the probability of a case being detected. We further estimate the number of undetected cases conditional upon the outbreak still being live, the epidemiological parameter R_0_, and the date on which the spillover event itself occurred. Our method requires minimal data to be effective. While our calculations were performed after the event, the real-time application of our method has potential value for public health responses to cases of emerging viral infection.

## Introduction

Viral transmission from animals to humans poses a serious threat in terms of its potential to generate novel pandemics[1]. Influenza viruses have a track record of causing serious impacts upon human health, with the 1918, 1957, and 1968 pandemics each being associated with more than one million deaths[2,3].

While such pandemics are rare events, they occur in a context of much more frequent animal-to-human spillover events: In 2022, epidemiological monitoring detected close to 60 cases of avian and swine influenza infection in humans worldwide[4]. While most of these events do not lead to large numbers of people being infected, at the very earliest stages of detection the distinction between an outbreak that will remain localised, and an outbreak that will go on to seed a pandemic, may be small. Efforts are required to understand spillover events in their earliest stages.

Statistical approaches for understanding outbreaks work on very different quantities of data. The SARS-CoV-2 pandemic saw the implementation of a broad range of computational and statistical approaches to track the nature and impact of the virus. Studies early in the pandemic characterised the epidemiological properties of the virus[5,6], and produced estimates of the epidemiological reproductive number R_0_ in different contexts[7]. Methods for ‘nowcasting’ combined multiple datasets to estimate local levels of viral prevalence[8,9]. A UK-wide project generated and shared hundreds of thousands of SARS-CoV-2 viral sequences[10]. Genome sequence data were used to study virus evolution and transmission on multiple scales[11–13].

Soon after a spillover occurs, data limitations may present themselves. Where multiple instances of infection from an emerging pathogen are observed, epidemiological models can highlight possible viral adaptation [14]. Inferences can be made of the epidemiological transmission parameter R_0_[15], and of differences in R_0_ across settings[16]. Sequence data can be used to assess the evolutionary origins and epidemiological characteristics of emergent viruses[17,18].

At the very earliest stages of an outbreak, data limitations may be severe; the detection of a spillover event may begin with a single case of infection. To explore what might be achieved in this minimal case, we here investigate a case of human influenza A(H1N2)v virus infection, observed the UK in November 2023. The detected case had no known contact with pigs, so was here assumed not to be the first case in the outbreak[19]. After the one detection, no further observations of infection were made. Considering the period immediately following the detection, we use a novel method to estimate the probability of the outbreak having ceased, and the potential number of undetected cases of infection. We discuss the potential for minimal datasets to inform the public health response in the days following the detection of a viral spillover event.

## Results

As a precursor to our method, we estimated the probability of a single case of influenza A(H1N2)v being detected as being between 4% and 10%. Publicly available data shows that in week 48 of 2023 the UK rate of GP consultations for influenza-like illness (ILI) was 4.6 per 100,000 individuals, equal to a national total of approximately 3,100 consultations per week.

Following these consultations, 557 samples were tested as part of a sentinel swabbing scheme, suggesting that approximately 18% of GP consultations for ILI led to swabbing and further testing[20]. Following the detection of a case, swabbing was increased in the Yorkshire and Humber region, where the case was found[20]. While data specific to the UK population are sparse, published estimates suggest that between 25 and 50% of individuals with ILI might seek healthcare[21,22]. Our estimate range was derived from these values.

We evaluated the properties of the A(H1N2)v influenza outbreak using likelihood free rejection sampling, first carrying out a form of historical nowcasting. Given a day following the detection of the outbreak, and supposing the knowledge that no further cases had been detected up until that day, we generated large numbers of simulated influenza outbreaks, identifying the statistical properties of simulations that matched the observed data up until the considered day. In this manner we estimated that, fourteen days after the first detection of A(H1N2)v influenza, the probability that the outbreak had died out was between 66% and 88% (Figure 1A). The range in this value reflects uncertainty in the probability of a case being detected, with higher detection probabilities leading to greater probabilities of the outbreak having ended. At a detection rate of 4%, we inferred that the outbreak could be said to have ended with 95% confidence by day 29 after the first detection; the same conclusion could be drawn by day 19 given a detection rate of 10%.

**Figure 1:**
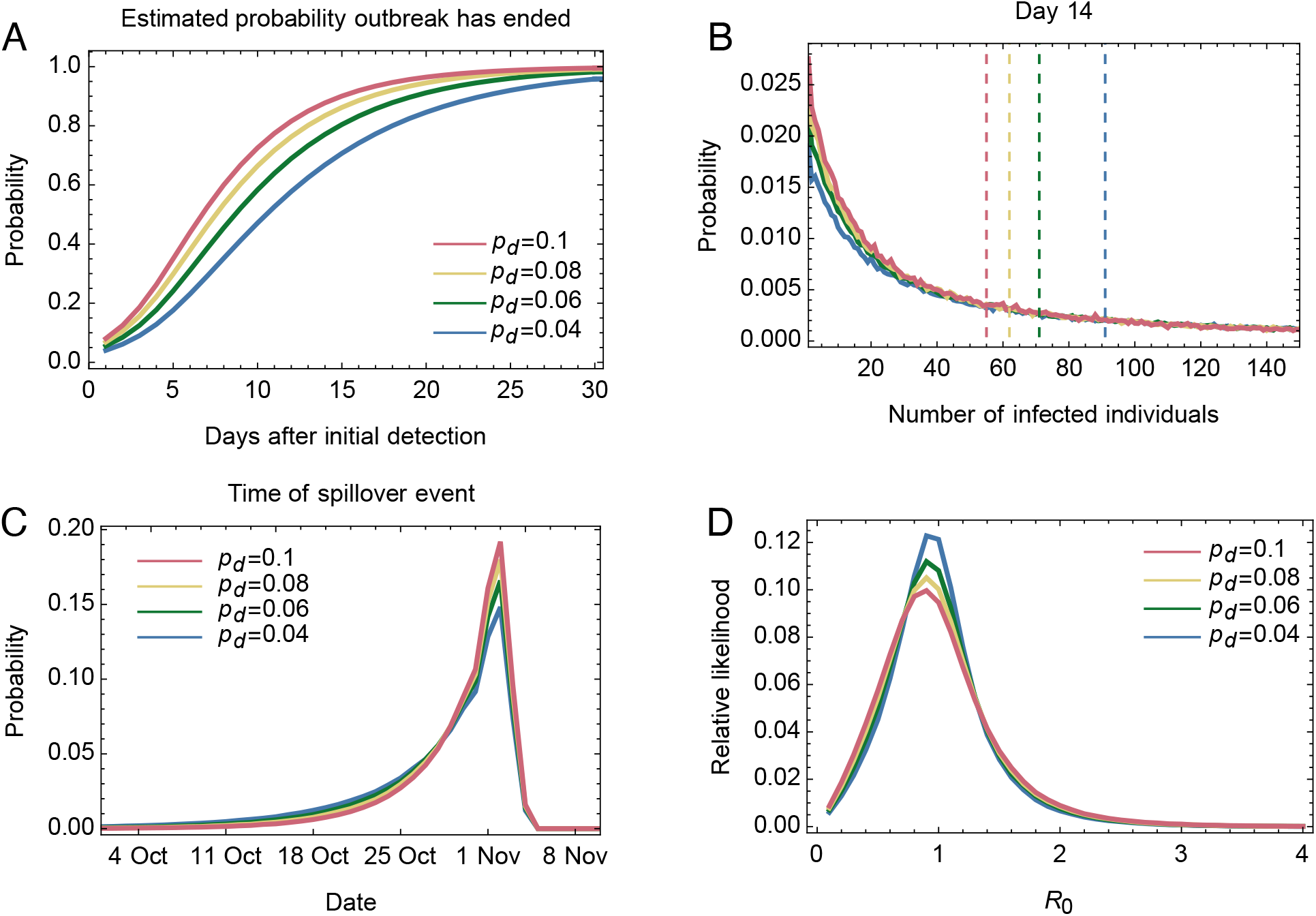
Sta2s2cs describing the influenza A/H1N2 spillover event calculated using our bootstrapping method. **A**. Time-dependent probability that the outbreak had died out. The value p_d_ describes the probability of a case of A/H1N2 influenza being detected. **B**. Calculated distribution describing the time at which the first case in the outbreak was infected. The detection of the first case was on 23^rd^ November. **C**. Calculated distribution of the number of infected individuals 14 days aQer the date of the first detection, conditional on the outbreak having not died out. The vertical dashed lines show the median values of each distribution. **D**. Estimate of the epidemiological parameter R_0_ for the influenza AtiH1N2 virus involved in this spillover event.

We further estimated the number of active cases of infection, conditional upon the outbreak having not died out. In the considered case, the median number of active cases fourteen days after the detection of the first case was estimated as between 55 and 91 (Figure 1B). Estimated numbers of active cases decreased slowly with time, considering days successively further from that of the detection (Supplementary Figure 1). Our model therefore suggested a dichotomy of potential circumstances: The outbreak was likely to have died out, but could have involved numbers of active cases if it was still live. We note that, soon after the detection of the first case, there exists the potential for the outbreak to involve very large numbers of cases. This result is explained by the 18-day delay between symptom onset and detection for the first detected case[23]. Given a high R_0_, this delay would have been sufficient for the virus to have spread substantially within the population. More rapid identification of outbreaks limits the potential for spread prior to detection.

We made a retrospective estimate of the time of the first, undetected, case of infection; estimates were made a nominal date 90 days after the first detection. Our model predicted that this first infection occurred 22 days prior to the first detection, on 1^st^ November, only a few days before the first detected case became symptomatic (Figure 1B). The inferred likelihood function is steeply skewed, with the mean of this distribution between 25 and 28 days prior to the first detection, varying with the detection rate. However, the interval between the time of the first infection and the time at which the first detected individual became symptomatic was likely short, with the detected individual potentially being a direct contact of the index case.

We also made a retrospective estimate of R_0_ for this outbreak, obtaining a value of 0.9 (Figure 1D). This lies below the threshold necessary to sustain an outbreak, and is lower than estimates for seasonal influenza viruses, for which R_0_ is in the region of 1.3[24]. The very sparse data used in making our estimate led to a large degree of uncertainty in this estimate; further data would sharpen the inferred distribution.

Exploring our data further, we examined the extent to which our retrospective estimates of R_0_ and the time of first infection could have been made closer to the observation of the first case. In the first few days after the detection of the case there was little power to rule out large values of R_0_ (Supplementary Figure 2A), but large values were progressively excluded with time. By contrast the inferred time of the first case of infection was relatively stable, with estimates calculated a few days after the observation being very close to our final estimate (Supplementary Figure 2B).

Our results are dependent upon the prior distribution chosen for R_0_. By default we used a uniform prior between zero and 4, but the highest values in this range are beyond those previously observed even for the 1918 pandemic virus[25]. Recalculating results with a uniform prior between zero and 2 did not strongly affect our results (Supplementary Figure 3).

## Discussion

Examining data describing a spillover event of a swine influenza virus into the human population we used a method of rejection sampling to explore what can be learnt, both at the time of the immediate response, and in retrospect, from the limited public data describing this event. Our method provided time-dependent estimates of the probability that the outbreak had died out, and for the number of undetected cases in the case that the outbreak was still live. It further provided estimates for the date of the spillover event itself, and for the parameter R_0_. Our model achieves what it does because both the observation of a case of infection, and the subsequent non-observation of cases, are informative for the model: The failure to observe a second case of infection is an important piece of data.

Our work pushes at the boundary of the amount of data required to learn about a spillover event, showing that even minimal data are sufficient to draw early and provisional conclusions about the outbreak. Our approach may be of value in a public health context: Estimating the certainty with which we can say whether an outbreak is likely to have died out could inform decisions about the investigation of cases and the resources applied to this task.

Simultaneously, the provisional nature of our conclusions should not be underemphasised.

Methods such as contact tracing would provide substantially more information about the potential for the virus to have spread, while genome sequencing of any further cases would facilitate phylogenetic and other genomic approaches to epidemiology.

Our approach is limited by its simplicity. For example, we assumed a homogeneous population, in which each infected person is equally infectious to the others, and neglected the potential for evolution to change the infectivity of the virus. We modelled data collection in a simple manner, assuming for example a fixed time between symptom onset and test result. We note that the epidemiological dynamics of infection, expressed as distributions of the time to symptom onset and to infecting others, are of critical importance to our method, but were of necessity based upon distributions inferred for other influenza strains: The A/H1N2 virus detected potentially would not mirror seasonal influenza in this way. Many of the assumptions made by our method could be elaborated upon, for example by allowing for heterogeneity in transmission[26]. The limited data available in this case did not encourage the use of more complex models.

The variation in our results under different detection scenarios highlights the potential value of improved systems for detecting cases of infection following a spillover event. In this case, detection efforts were stepped up in the region of the detection of the first case. More thorough and faster testing provides a greater certainty that a spillover has not led to a persistent outbreak and reduces the potential for an outbreak to grow undetected.

While we have applied our model to a specific event, describing the spillover of influenza A(H1N2)v into the human population in the UK, our approach has the potential for broader use. Given reasonable estimates for epidemiological parameters, viruses other than influenza could also be modelled. Events involving more than one detection of a positive case could also be assessed, though as the number of cases of infection increases our approach becomes less computationally efficient. Once large amounts of data become available, alternative methods for epidemiological inference are likely to perform better than our own. Our approach is of potential value in the first stages of an outbreak when data are most limited.

## Methods

We used likelihood free rejection sampling to evaluate the likely state of the outbreak underlying the available data. This method generated a large number of simulated outbreaks, retaining only those which were compatible with the data collected from the A(H1N2)v outbreak, counting the number of cases detected on each day, and assuming that a given number of days, denoted by t_o_, had passed since the day on which the first detected case was observed. For the historical now-casting calculations simulations which exactly matched the data up to time t_o_ were accepted. The R_0_ parameters of the accepted simulations provided samples of the posterior distribution of this statistic. The properties of these simulations for different values of t_o_ were used to estimate the properties of the outbreak. Where retrospective estimates of parameters were made, these were calculated at the point t_o_ = 90 days.

### Simulation of outbreaks

Time was modelled discretely, in units of whole days. Each simulated outbreak started with a primary case. The simulation then proceeded day by day. Infected individuals were assumed to become symptomatic a random number of days t_s_ after being infected. Every symptomatic individual then infected a total of R others, with the time of each infection event occurring a random number of days t_i_ after symptom onset, and where R was Poisson distributed with parameter R_0_. The random numbers t_s_ and t_i_ were drawn from Weibull distributions with parameters based upon published parameters describing influenza infection[27], but with samples rounded to the nearest integer. Specifically, where w is the cumulative distribution function

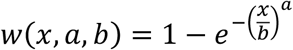

*t*_*s*_had probability mass function

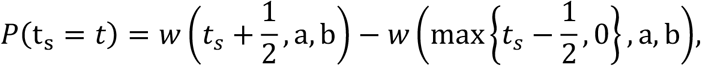

And similarly for t_i_. Parameters for the distribution of t_s_ were given by a=7.4026 and b=1.7375, while parameters for the distribution of t_i_ were given by a=1.0314 and b=1.0025.

For each infected individual in our simulation we randomly determined whether the case was detected. Detection was modelled as occurring with fixed probability p_d_. Detection was assumed to occur 18 days after the day of symptom onset, following data from the influenza A(H1N2)v case[23]. The exception to this rule was the primary case in the outbreak. Following information that the index case had no contact with animals, the primary case was assumed not to have been detected[19].

Each simulated outbreak was continued until it either died out, with no more cases of infection existing, or until the first day we were certain whether or not the numbers of cases detected in the simulation matched the number of detected cases in the dataset up to the observation time.

We assumed a uniform prior over the epidemiological parameter R_0_ within the window [0.1, 4.0]. Rather than randomly sample, we performed a grid search, conducting 10^6^ simulations for each of 40 equally spaced discrete values of R_0_ from 0.1 to 4.0.

Once simulations were complete, we calculated the proportion of simulations for each R_0_ that were accepted, 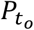 (accepted|*R*_0_). Normalising this statistic gave the posterior probability of R_0_ at a given observation time t_o_,

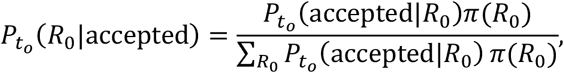

where the prior *π*(*R*_0_) is uniform. Using the accepted simulations, we estimated properties of the viral population, using the formula

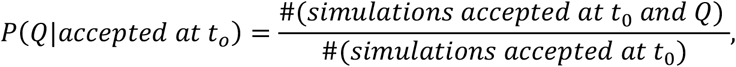

for properties Q including the number of infected individuals being k at time t_i_ days after the detected case, the outbreak having died out at time t_i_ days after the detected case, or the time of the first infection having occurred a specific number of days before the detected case.

In the calculations above, we assumed that infection lasted for seven days following infection. We note that alternative prior distributions for R_0_ could be used in our calculation; we show in Supplementary Information the results of placing a lower upper bound on this statistic.

## Data/Code availability

All data used for this analysis was obtained from publicly available sources. Our code is named OINK (Outbreak Inference given Negligible Knowledge) and is available from https://github.com/cjri/OINK/.

## Acknowledgements

We acknowledge funding from the UK Medical Research Council (grant number MC_UU_00034/6). We thank Margaret Hosie and Antonia Ho for comments on an earlier version of this mansuscript. We thank the Pandemic Preparedness team at the MRC University of Glasgow Centre for Virus Research, and Dr Robert Goudie for discussions.

