## SupplementaryFigures for "Epidemiological inference at the threshold of data availability: an influenza A(H1N2)v spillover event in the United Kingdom"

**Supplementary Figures**

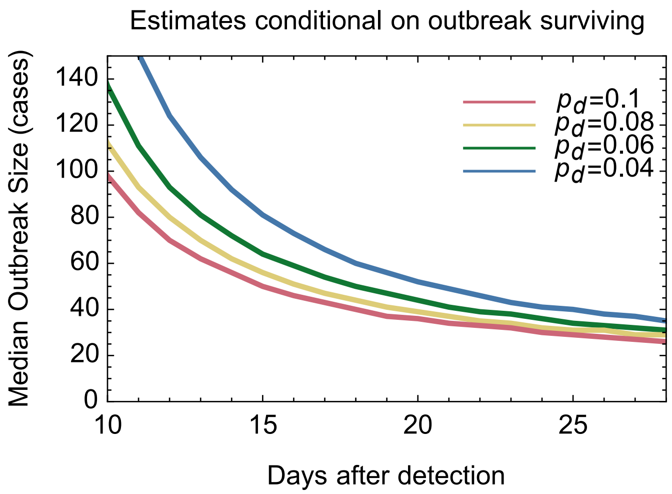


**Supplementary Figure S1:** The median number of cases in the outbreak, conditional on the outbreak not having died out, estimated at different number of days having passed after the initial detection.

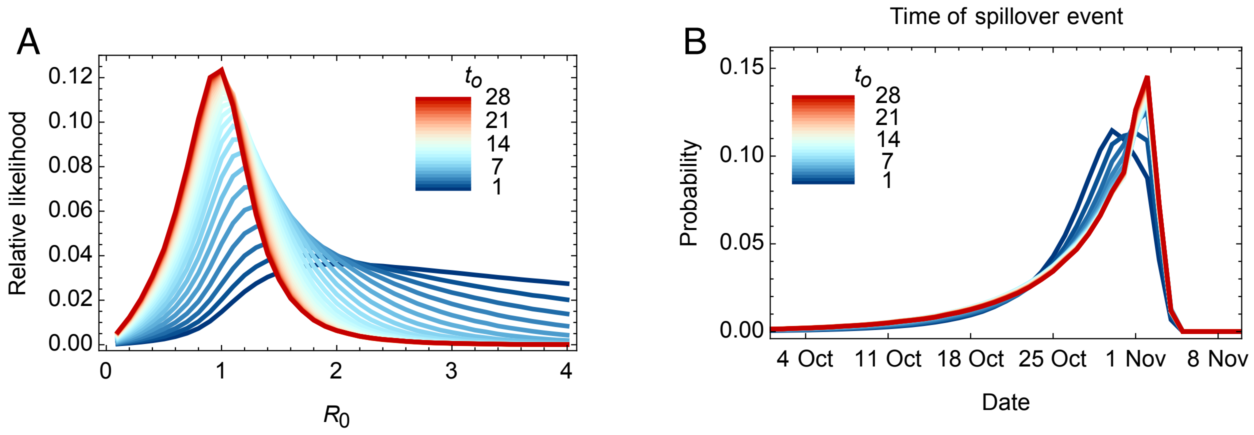


**Supplementary Figure S2: Variation between distributions of parameters inferred at times shortly after the detection of the first case at $t_o$=0.** Data are shown for the detection probability $p_d$=0.04. **A.** The time-dependent posterior distribution of $R_0$, calculated under the assumption that no further cases are detected after the first case of infection. **B.** The distribution of the time at which the first case in the outbreak was infected.

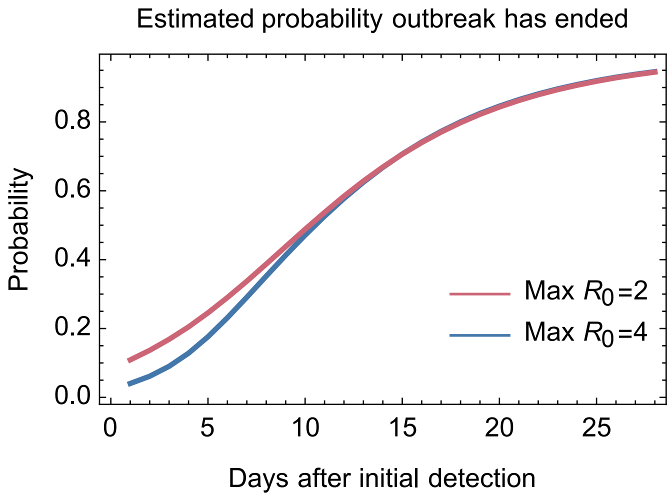


**Supplementary Figure S3: Dependence of the inferred probability that the outbreak had ended upon the upper limit of the prior distribution of $R_0$.** Data are shown for the detection probability $p_d$=0.04. In both cases a uniform prior for $R_0$ was applied.
